# Precapillary sphincters control cerebral blood flow

**DOI:** 10.1101/657486

**Authors:** Søren Grubb, Changsi Cai, Bjørn O. Hald, Lila Khennouf, Jonas Fordsmann, Reena Murmu, Aske G. K. Jensen, Stefan Zambach, Martin Lauritzen

## Abstract

Active nerve cells produce and release vasodilators that increase their energy supply by dilating local blood vessels, a mechanism termed neurovascular coupling, which is the basis of the BOLD (blood-oxygen-level-dependent) functional neuroimaging signals. We here reveal a unique mechanism for cerebral blood flow control, a precapillary sphincter at the transition between the penetrating arteriole and the first capillary that links blood flow in capillaries to the arteriolar inflow. Large NG2-positive cells, containing smooth muscle actin, encircle the sphincters and rises in nerve cell activity cause astrocyte and neuronal Ca^2+^ rises that correlate to dilation and shortening of the sphincter concomitant with substantial increases in the RBC flux. Global ischemia and cortical spreading depolarization constrict sphincters and cause vascular trapping of blood cells. These results reveal precapillary sphincters as bottlenecks for brain capillary blood flow.

## Introduction

Neurovascular coupling (NVC) is the signalling mechanism that links neuronal activity to local increases in cerebral blood flow^1^. Ca^2+^ rises in neurons and astrocytes trigger release of vasoactive compounds that dilate capillaries and penetrating arterioles, which in turn increases local blood flow. The activity-induced rises in blood flow are based on coordinated changes in the diameters of arterioles and capillaries which in turn are regulated by Ca^2+^ fluctuations within the vascular smooth muscle that circumscribe arterioles and the pericytes which ensheath capillaries close to the penetrating arteriole^2–5^. Intracortical arterioles branch into the capillary networks that supply the cortical layers with oxygen and glucose^6^. It is unclear, however, how the organization of blood supply can ensure a roughly equal perfusion of capillary networks at different cortical depths. The organization encounters two competing obligations: preservation of perfusion pressure in the penetrating arteriole along the entire cortical depth, which is essential for adequate blood flow to all layers, yet the brain tissue must be shielded from the mechanical impact of blood pressure rises. Here, we reveal the structure and function of brain precapillary sphincters, which may serve exactly these two purposes: capillary protection from systolic pressure spikes and preservation of perfusion pressure despite capillary branching from the penetrating arteriole. We characterized precapillary sphincters as mural cells encircling an indentation of blood vessels exactly where capillaries branch off from penetrating arterioles. The sphincter cells had a bulbous soma similar to brain pericytes, contained α-smooth muscle actin and were ensheathed by structural proteins. Precapillary sphincters were present at most but not all proximal capillary branches of penetrating arterioles (PA) and with a decrease in occurrence from upper to lower cortical layers, an ideal position to facilitate a balanced perfusion pressure along the PA and for brain protection against arterial pressure pulsations. While precapillary sphincters have been known for almost a century^7^, their existence in all vascular beds except for the mesentery^8–10^ remains controversial^11,12^. This paper provides unequivocal structural and functional evidence for brain precapillary sphincters and examines their role in neurovascular coupling and in pathology during cortical spreading depolarization (CSD) and global ischemia following cardiac arrest.

## Results

### Precapillary sphincters mainly locate to the proximal branch-points of penetrating arterioles

We identified precapillary sphincters in mice expressing dsRed under the NG2 promoter as lobular dsRed-positive cells encircling an indentation of the vessel lumen at PA branch-points (Fig. 1a). Precapillary sphincters were most often followed by a distention of the lumen, which we denoted “the bulb”. The dsRed signal from the precapillary sphincter was usually brighter than the dsRed signals from other mural cells on the PAs and 1^st^ order capillaries indicating high NG2-expression. However, the dsRed signal from the bulb region was low compared to the rest of the 1^st^ order capillary (Fig. 1a,b,d), which suggested low pericyte coverage. We could show that precapillary sphincters and bulbs were not only present in anesthetized mice, but also in awake mice with chronic cranial windows *in vivo* (Fig. 1c and Extended data Fig. 3, n = 4) and in anesthetized NG2-dsRed mice with thinned skull over the barrel cortex *in vivo* (Fig. 1b, Extended data Fig. 2 and Supplementary video 1, n = 3). *Ex vivo* studies revealed that the NG2-positive cells encircling the precapillary sphincter were individual cells encompassing the sphincter at the branchpoint and not processes of mural cells extending from the PA (Fig. 1d). DAPI stain of coronal brain slices, revealed that penetrating arterioles were covered with smooth muscle/pericyte hybrids (Fig. 1e), indicating a continuum of mural cell cyto-architecture from pial arterioles to 3^rd^ order capillaries as previously described^13–15^.

**Figure 1.**
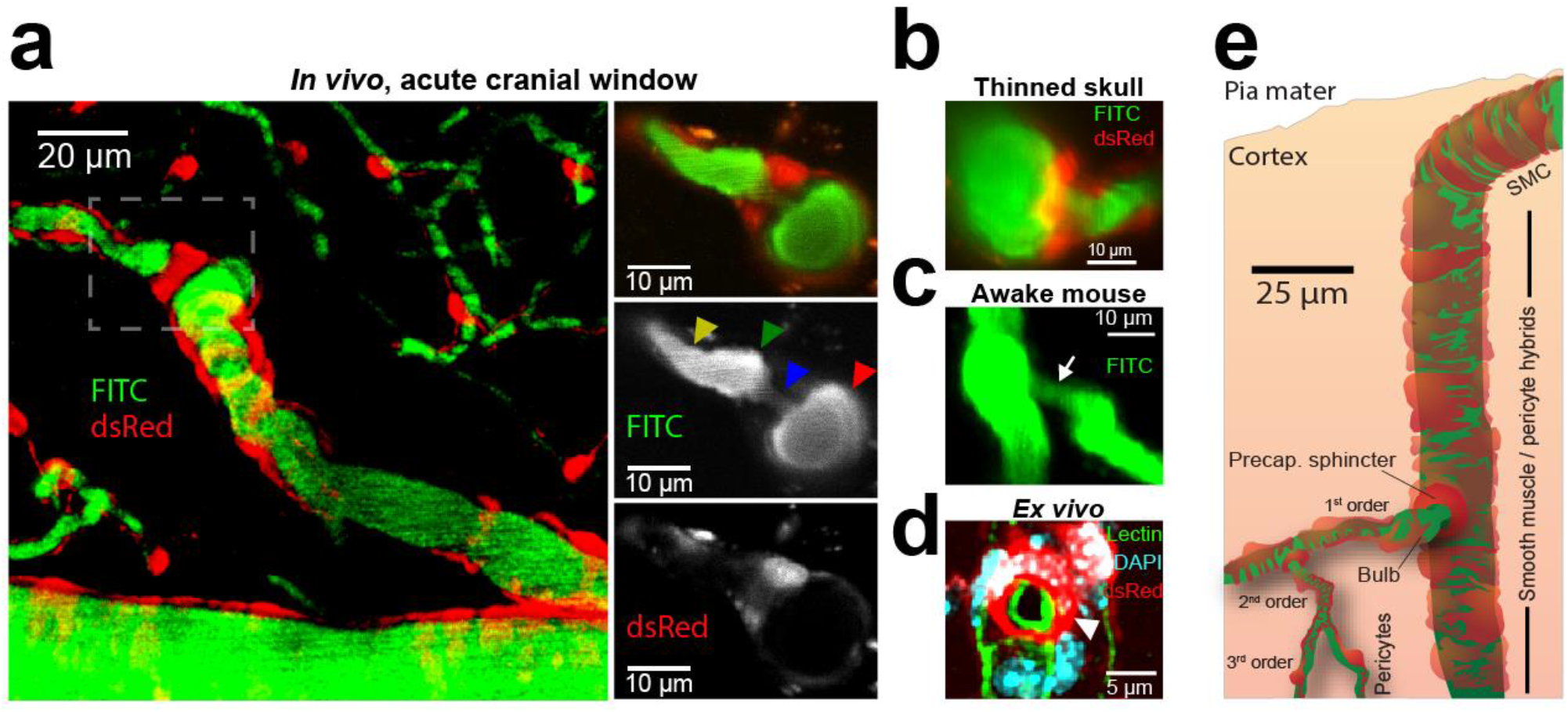
Precapillary sphincters are found on the proximal branches of penetrating arterioles. **a**, *Left panel:* Maximal intensity projected *in vivo* two-photon laser scanning microscopy image of an NG2-dsRed mouse barrel cortex. An indentation of capillary lumen is observed at the branching of the penetrating arteriole (PA)and is encircled by bright dsRed cell(s) (dashed insert). This structure is denoted a precapillary sphincter. Immediately after the sphincter a sparsely dsRed-labeled distention of the capillary lumen is observed, which we refer to as “the bulb”. *Right panels:* Single z-plane showing overlay, FITC-channel, and dsRed channel of the dashed insert. Arrows indicate the PA (red), sphincter (blue), bulb (green) and 1^st^ order capillary (yellow). **b-d**, Local TPLSM projections of precapillary sphincters in cortex of: **b**, a thinned skull mouse *in vivo*, c, an awake mouse harboring a chronic cranial window *in vivo.* White arrows mark the precapillary sphincter, and **d**: an *ex vivo* coronal slice of a FITC-conjugated lectin (green) stained NG2-dsRed mouse (red) with DAPI staining (blue) of nuclei. The precapillary sphincter cell nucleus is arched, as it follows the cell shape. **e**, Schematic of a PA with a precapillary sphincter at the proximal branch-point based on *ex vivo* data. The morphology and location of NG2-dsRed positive cells on the vascular tree are indicated.

Having established the structure of precapillary sphincters, we examined their occurrence and localization within the cortical vascular network. In keeping with the work of Duvernoy et al.^6^, we identified a range of PA subtypes (Fig. 2b) that differed in size, branching pattern and cortical penetration. Precapillary sphincters were predominantly localized to the upper layers of the cortex (Fig. 2c) and were mainly observed at proximal PA branch-points (Fig. 2d) of relatively large PAs branching into relatively large 1^st^ order capillaries (Fig. 2e,f). As larger proximal vessels carry higher blood pressures than downstream vessels, this localization indicated that sphincters contributed to pressure distribution. The bulb usually succeeded a sphincter, but was less prevalent and did not display the same positive correlation with the diameter of the 1^st^ order capillary as the sphincter (Fig. 2e), i.e. bulbs were observed when the PA diameter was large compared to the 1^st^ order capillary (Fig. 2f). For branches positive for a precapillary sphincter, the average diameter of the PA was 11.4 ± 0.6 μm, the precapillary sphincter was 3.4 ± 0.2 μm, the bulb was 5.8 ± 0.2 μm, and the 1^st^ order capillary was 5.3 ± 0.2 μm (mean ± SEM). As per Poiseuille’s law (adjusted for flow velocity, Fig. 2g), a lumen diameter around 3-4 μm is precisely at the border of very high flow resistance, providing an effective means of changing the pressure drop per unit length. We conclude that precapillary sphincter complexes (sphincter *and* bulb) 1) are characterized by an indentation of lumen at the branch-point encircled by a sphincter cell usually followed by a distention (the bulb) and 2) are common at proximal PA branchpoints, predominantly at larger PAs in the mouse cortex.

**Figure 2.**
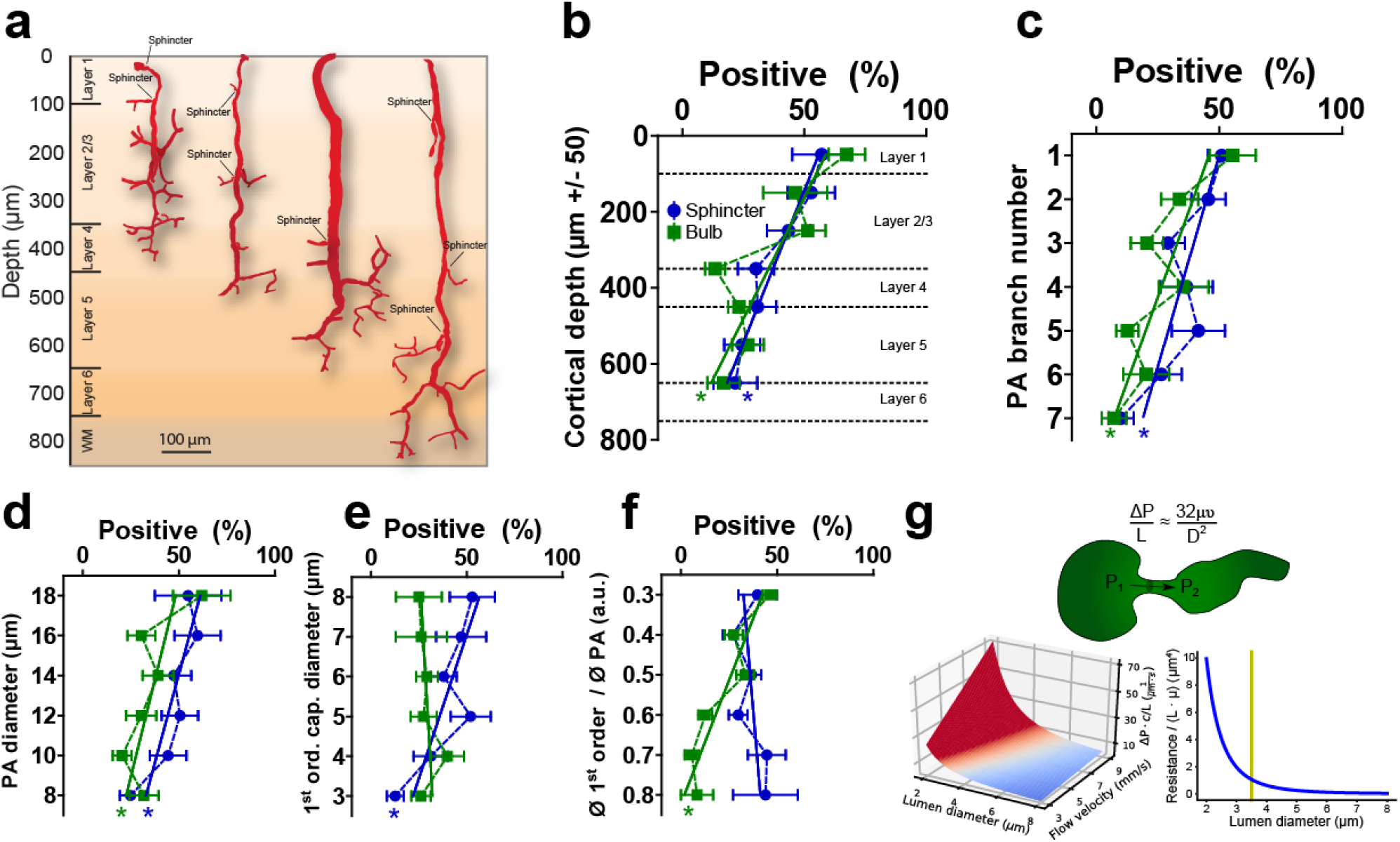
Presence, location, and biophysics of cortical precapillary sphincters suggest a function in pressure equalization across the PA. **a**, Representatives of four PA subtypes^6^ reaching different cortical layers based on *ex vivo* data. Precapillary sphincters are found at varying depths. b-f, Dependency of the presence and location of precapillary sphincters and bulbs (binned quantification) on various parameters. Criteria of a positive presence of sphincter or bulb at a branch point: sphincter <0.8 and bulb >1.25 times the diameter of a 1^st^ order capillary, N = 6-8 mice, ±SEM, linear regression, * = slope deviates significantly from 0. **b**, dependency on cortical depth (bin size 100 μm). **c**, dependency on PA branch number (counting from the proximal end). **d**, dependency on PA diameter (bin size 2 μm). **e**, dependency on 1^st^ order capillary diameter (bin size 1 μm). **f**, dependency on 1^st^ order cap / PA diameter ratios (bin sizes as in d and e). **g**, *Top panel:* Illustration of a pressure drop across a precapillary sphincter and a modified expression of Poiseuille’s law (ΔP: pressure difference, L: unit length, μ: viscosity, υ: flow velocity). *Lower left:* Graphical illustration of Poiseuille’s law showing how the pressure difference times viscosity per unit length depends on the cylindrical lumen diameter and flow velocity. Note how the pressure difference increases with lumen diameters below ~ 4μm. *Lower right:* An equivalent representation of how flow resistance per unit length and viscosity depends on lumen diameter according to Poiseuille’s law.

### Precapillary sphincters actively regulate capillary blood flow

Having established the occurrence and morphology of precapillary sphincter complexes, we examined their role in blood flow regulation. First, we confirmed expression of α-smooth muscle actin (α-SMA) within the precapillary sphincter cell in coronal slices of NG2-dsRed mice (Fig. 3a, vascular lumen and cell nuclei costained using lectin and DAPI, respectively. See also Extended data Fig. 4 and Supplementary video 2). Next, we analyzed the vasomotor responses of the PA, precapillary sphincter, bulb, and 1^st^ order capillary vessel segments in response to electrical whisker pad stimulation in an *in vivo* two-photon setup (Extended data Fig. 1). Careful placing of linear regions of interest (ROIs) in hyperstacks of two-photon images were used to avoid inter-segmental interference in diameter calculations before and during whisker stimulation (Fig. 3b-c and Supplementary video 3). Precapillary sphincters dilated during stimulation followed by a post-stimulus constriction, denoted the post-stimulus undershoot at 20-30 s after stimulation. Four-dimensional hyperstack imaging allowed us to confirm that the undershoot was not an artifact of drift in the z-axis. Relative diameter changes at the sphincter were significantly higher compared to the PA and the rest of the 1^st^ order capillary during both dilation (33.75±4.08%, Fig. 3e and Extended data Table 1) and undershoot (−12.40±2.10%, Fig. 3f and Extended data Table 1). To estimate the corresponding changes in flow resistance per unit length, we applied Poiseuille’s law at baseline, maximal dilation, and maximal undershoot (Fig. 3g-i). The flow resistance of the sphincter at rest was significantly higher compared to the other segments and decreased significantly more (65.9% decrease, Fig. 3h) during dilation as compared to all other segments (40.8% for the 1^st^ order capillary, Fig. 3h). During the post-stimulus undershoot, flow resistance increased by 80.2% at the sphincter (Fig. 3i), which highlights the sensitivity of flow resistance to sphincter constriction and underscores the strategic control of flow resistance at the sphincter due to the power law relationship between diameter and flow resistance (Fig. 2g). Moreover, we observed that the length of precapillary sphincters decreased during stimulation and increased during undershoot (Extended data Fig. 5). According to Poiseuille’s law, shortening of the sphincter decreases the absolute flow resistance across the precapillary sphincter complex and vice versa, i.e. further reducing the pressure drop across the sphincter during functional dilation and increasing the pressure drop during the undershoot. We examined intracellular Ca^2+^ dynamics in neuronal somas and astrocytic end-feet enwrapping the vessel segments (Extended data Fig. 6). Neuronal somas and astrocytic end-feet responded with increases in intracellular Ca^2+^ upon whisker pad stimulation and in the undershoot phase (Extended data Fig. 6b-e). The fraction of ROIs responding was similar during dilation and undershoot and independent of the location of the end-feet on the vascular three (Extended data Fig. 6f).

**Figure 3.**
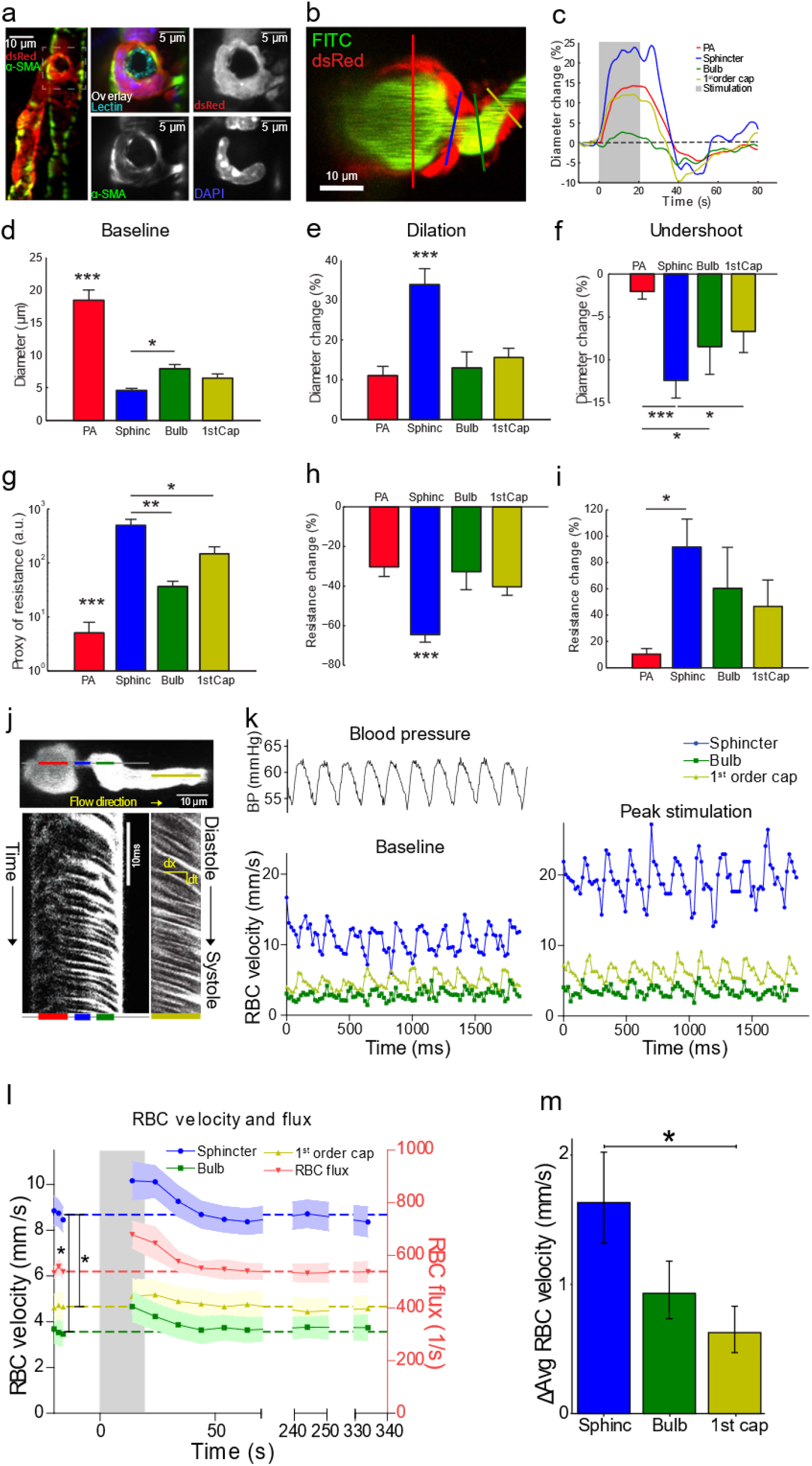
Precapillary sphincters are actively regulating blood flow upon functional stimulation. **a**, *Ex vivo* coronal slices of a FITC-lectin stained NG2-dsRed mouse immunostained for α-SMA. *Left panel:* Maximal projection of a PA with a precapillary sphincter at the 1^st^ order capillary branchpoint, marked area is shown in right panels. *Right panels*: Local maximal intensity projections of the precapillary sphincter region of either dsRed, α-SMA, DAPI, or all channels including FITC-lectin overlayed. **b-i**, *In vivo* whisker pad stimulation experiments (anesthetized NG2-dsRed mice) using maximal intensity projected 4D data obtained by two-photon microscopy. **b**, Maximal intensity projection of a PA branchpoint where the colored lines indicate the ROIs for diameter measurements for the vessel segments PA (red), precapillary sphincter (blue), bulb (green) and 1. order capillary (yellow). **c**, Representative time series of relative diameter dynamics in each vessel segment upon 20s of 5 Hz whisker pad stimulation (grey bar, start at time zero). **d**, Summary of baseline diameters (absolute values). **e**, Summary of peak diameter change upon whisker pad stimulation. **f**, Summary of peak undershoot phase after whisker pad stimulation. **g**, A proxy of flow resistance at baseline estimated using Poiseuille’s law. **h**, Relative change in flow resistance at peak dilation during stimulation. **i**, Relative change in flow resistance during the post-stimulation undershoot. **j-m**, RBC velocity and flux estimation. **j**, Resonance scanning allows for rapid repetitive line-scans in a single z-plane (upper panel). In the resulting space-time maps (lower panel), individual cells display in black with an angle proportional to the cell velocity. Red, blue, green and yellow lines indicate the regions of the line-scans deriving from the PA, sphincter, bulb, and the 1^st^ order capillary (1^st^ order capillaries were mostly scanned in consecutive experiment). **k**, Fluctuations in femoral artery blood pressure (left upper panel) and RBC velocity (left lower panel) correlated. During whisker pad stimulation (right panel), RBC velocity increased. **l**, Time series of RBC velocities and flux during whisker pad stimulation. RBC velocity at the precapillary sphincter was significantly higher than the bulb and 1^st^ order capillary at baseline and peaked around 10s after stimulation before returning to baseline. **m**, Summary of the difference between maximal and baseline RBC velocity during whisker stimulation. The Kruskal-Wallis test was used in d, g, and i to reveal differences among vessel segments followed by a Wilcoxon rank-sum test (with Holm’s p-value adjustment) for pairwise comparisons. LME models were used in e, f, h, and m to test for difference among segments followed by Tukey post hoc tests for pairwise comparisons. In m, the LME analysis was performed on log-transformed data to ensure homoskedasticity.

We next examined the correlation between red blood cell flux and diameter changes in response to whisker pad stimulation (Fig. 3j-m). RBC velocity fluctuated in synchrony with systolic and diastolic oscillations in arterial blood pressure (Fig. 3j,k). At rest, the average RBC velocity through precapillary sphincters was 8.7±0.6 mm/s (Fig 3l), significantly higher than for the bulb (3.6±0.6 mm/s) and the 1^st^ order capillary (4.7±0.6 mm/s), but correlated to the relative differences in resting diameters of the vessel segments. As shown in Fig. 2G, high RBC velocity through the narrow lumen of the precapillary sphincter amplifies the reduction of pressure across the sphincter due to high shear and thereby contributes to the protection of downstream capillaries from high pressures in large proximal PAs. From the baseline measures, the pressure drop per unit length is 4 times larger in the sphincter compared to the 1^st^ order capillary, assuming that RBC velocity and fluid velocity are roughly equal, see Fig. 2g. During whisker stimulation (Fig. 3l), both diameter and RBC velocity increased in each segment but significantly more at the precapillary sphincter than the 1^st^ order capillary (Fig. 3m). The flux of RBC’s through the precapillary sphincter complex increased by 25% from baseline to peak stimulation (mean flux rose from 543±25 to 679±50 cells per second, Fig. 3l). Given the high RBC velocity, a baseline flux in the first order capillary of around 550 cells per second is not surprising (see Extended data calculation 1). The sphincter, however, retained a dampening effect on pressure during peak stimulation, where the pressure drop per unit length was 3 times larger at the sphincter compared to the 1^st^ order capillary. RBC velocity and flux returned to baseline at 20-30s after end of stimulation (Fig. 3l), concurrent with the post-stimulus undershoot (Fig. 3c,f) ^17^.^16^ Before, during, and after whisker stimulation, we observed fluxes of single RBCs through the precapillary sphincter into the 1^st^ order capillary, which may optimize oxygen delivery to the tissue (Supplementary video 5). Collectively, our data suggest that the sphincter complex 1) protects downstream capillaries from blood pressure peaks in the proximal PAs, 2) actively regulates local diameter and RBC flux during functional stimulation, and 3) equalizes the distribution of RBCs entering the upper and lower cortical layers.

### Passive structural elements around the precapillary sphincter support the bottleneck function

The presence of a contractile sphincter-encircling pericyte, supports the notion of active regulation of the diameter at the precapillary sphincter. The indentation of the sphincter, however, might also be supported by passive elements to optimize the force-length relationship^17^. We therefore investigated whether passive structural elements constrained dilation at the sphincter by injecting papaverine (10 mM), a strong vasodilator, close to the sphincter (Fig. 4a-c). Papaverine blocks the contractility of the vascular smooth muscle cells, and by inference pericytes, by inhibiting vascular phosphodiesterases^18^ and calcium channels^19^. Under these conditions, passive structural elements of the vessel become the main factors that stabilize the vessel wall in face of the unaffected transmural pressure. Both before and after papaverine injection, the lumen diameter of the sphincter was significantly smaller than for the bulb and 1^st^ order capillary (Fig. 4c). Yet, the sphincter showed a significantly larger dilation in absolute and relative terms, as compared to the 1^st^ order capillary. Structural evidence of passive connective tissue was established by staining coronal slices of NG2-dsRed mice with either a collagen α1 type I (COL1A1) antibody or Alexa633 hydrazide, a marker of elastin^20^. Elastin was observed in the tunica intima of penetrating arterioles and at the precapillary sphincter, but not at the level of 1^st^ order capillaries (Fig. 4d). Collagen α1 type I staining was observed in the tunica externa of arterioles, precapillary sphincters, capillaries (Fig. 4e), and venules. Thus, common structural proteins ensheathed the precapillary sphincter. The data indicates that the active sphincter is supported by passive structural elements that assist the active pericyte in protecting soft brain tissue against pressure increases produced by dilation of the PA.

**Figure 4.**
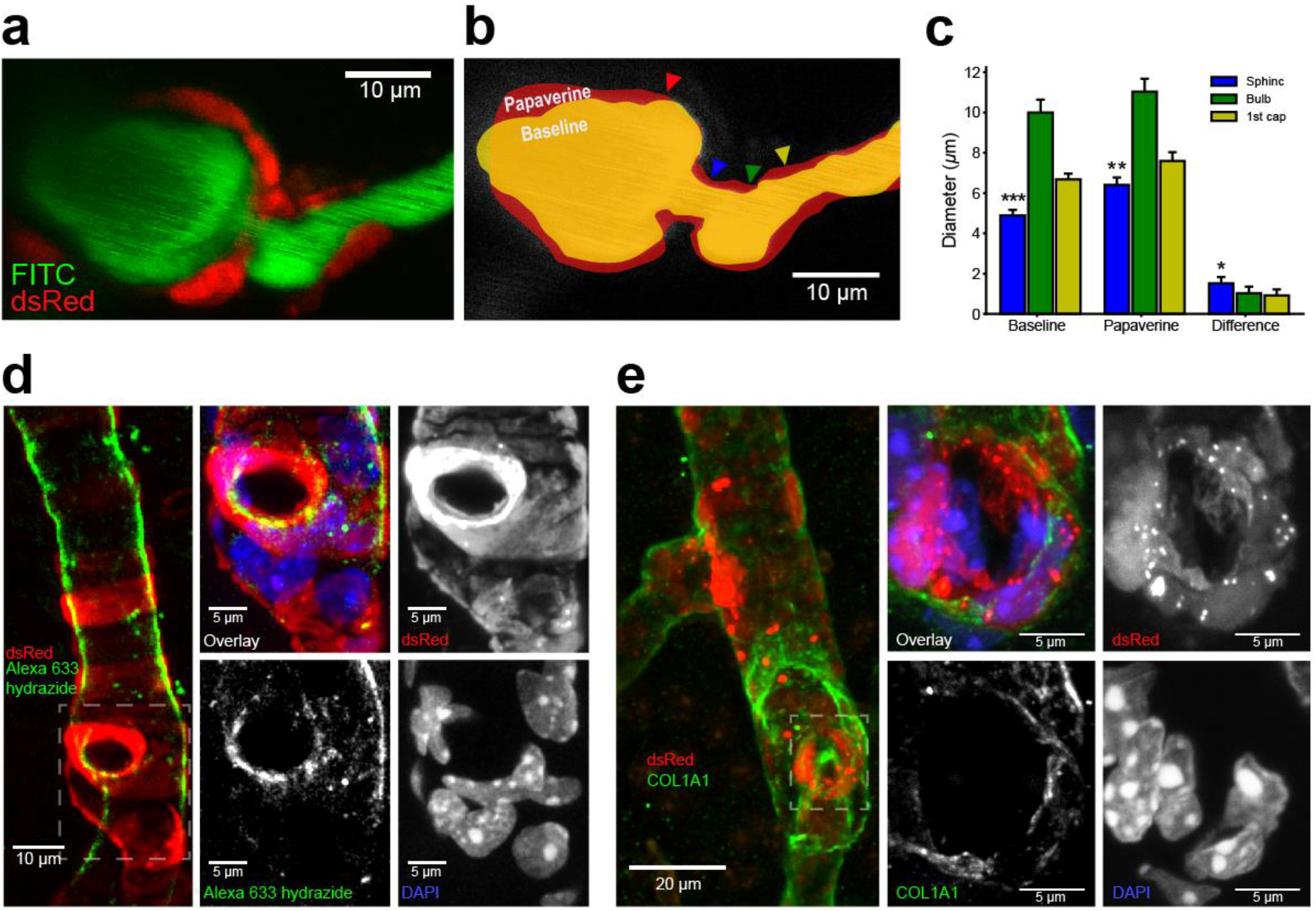
Precapillary sphincters harbor passive structural elements that delimit capacity for vasodilation. **a-d**, Papaverine (10 mM) was locally injected into the vicinity of precapillary sphincters to dilate the nearby vasculature. **a**, Representative maximal intensity projection of an NG2-dsRed mouse PA branch-point. **b**, Schematic showing the papaverine induced dilation (red) below an outline of the vessel lumen at baseline (yellow). ROI locations of individual vessel segments are marked by colored arrows. **c**, Absolute diameters of vessel segments 1) at baseline, 2) after papaverine addition, and 3) the difference before and after papaverine addition. The baseline dataset was analyzed using the Kruskal-Wallis test followed by a Wilcoxon rank-sum test (with Holm’s p-value adjustment) for pairwise comparisons. The papaverine and difference datasets were analyzed using LME models followed by Tukey post hoc tests for pairwise comparisons. **d**, Maximal intensity projections of coronal slices of NG2-dsRed mice stained with Alexa633 hydrazide and DAPI. *Left panel:* 20x magnification of a penetrating arteriole with a precapillary sphincter at the branchpoint. *Right panels:* 63x magnification of the precapillary sphincter and 1^st^ order capillary. Alexa633 hydrazide staining is strong at the sphincter but absent in the 1^st^ order capillary. **e**, Maximal intensity projections of coronal slices of NG2-dsRed mice stained with COL1A1 antibody and DAPI. *Left panel:* 20x magnification of a penetrating arteriole with two branches. *Right panels:* 63x magnification of the precapillary sphincter at the lower branch.

### The precapillary sphincter in cortical spreading depolarization and during global ischemia

In the healthy mice considered so far, precapillary sphincter complexes displayed an active role in regulation and protection of downstream capillary blood flow (Fig. 3,4). As observed for the undershoot (Fig. 3f,i), the flow resistance of the sphincter may increase strongly under physiological conditions (Fig. 2g). This observation prompted the question of whether sphincters constrict in brain pathology. Hence, we investigated sphincter dynamics during cortical spreading depolarization (CSD) waves that are caused by disrupted brain ion homeostasis and known to cause prolonged vasoconstriction^4^. Microinjection of 0.5 M potassium acetate in the posterior part of the somatosensory cortex elicited CSD that triggered a triphasic sequence of changes the in the diameter and flow of cortical blood vessels consisting of: (I) a brief initial constriction followed by (II) a longer-lasting dilation and (III) a prolonged vasoconstriction (Fig. 5a-c, supplementary video 6). Whereas the maximal constriction relative to baseline in phase I was similar among vessel segments, the maximal relative dilation of the precapillary sphincter in phase II was greater than for the bulb and the 1^st^ order capillary (Fig. 5d, 39±8%, vs. 22±3 and 21±4%) but not different from maximal dilation during whisker pad stimulation (34±4%, Fig. 3e) or local injection of papaverine (32±8%, Fig. 4c). During phase III, the precapillary sphincter constricted more (26.2%) than the PA and the bulb and generated a doubling in flow resistance (Fig. 5e). This was occasionally accompanied by transitory entrapment of RBCs at the sphincter that occluded the 1^st^ order capillary (Supplementary video 4), consistent with the high increase in flow resistance. These results show that the sphincter constricts markedly in CSD, which is likely to be highly important for the associated long lasting decreases in cortical blood flow that follows CSD^21^.

**Figure 5.**
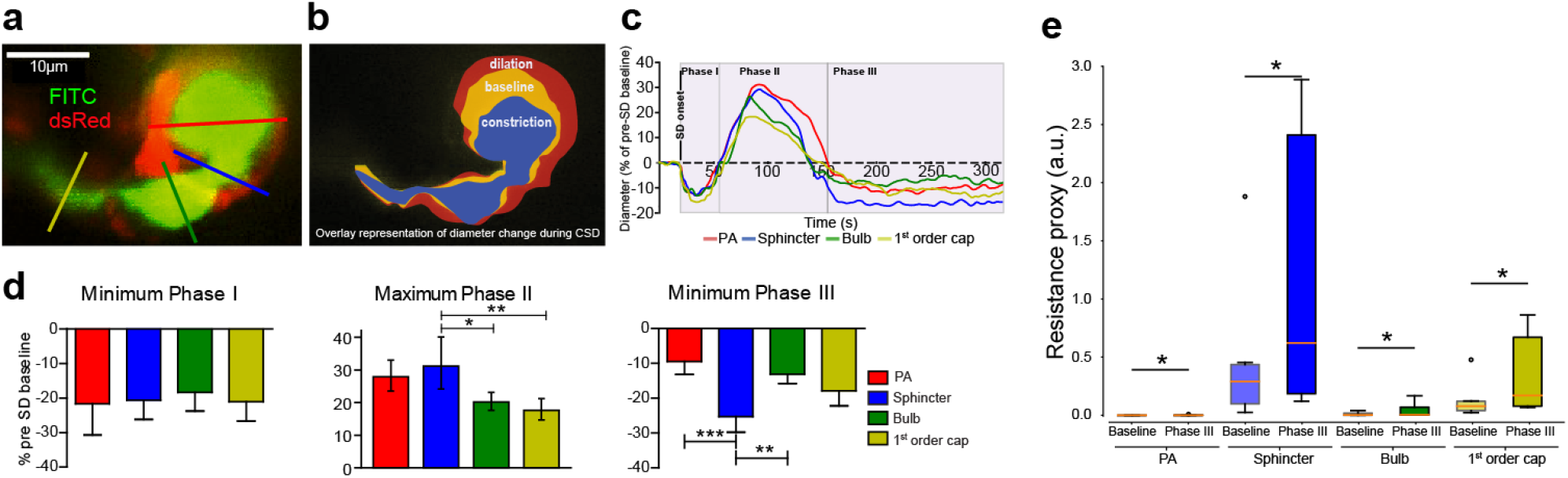
a-d, Precapillary sphincters are vulnerable to conditions of general constriction. Cortical spreading depolarization was elicited in the posterior part of the somatosensory cortex by microinjection of potassium acetate during imaging of the precapillary sphincter. **a**, Representative maximal intensity projection of a FITC-dextran loaded NG2-dsRed mouse at a precapillary sphincter. Colored lines mark the ROIs for diameter measures. **b**, Overlaid outlines of baseline (yellow), phase II dilation (red) and phase III constriction (blue). **c**, Representative time series of diameter changes within vessel segments during the three phases of CSD. **d**, Summaries of maximal diameter changes within vessel segments during phase I, II and III of the CSD. During phase II, the PA and sphincter dilated significantly more than the 1^st^ order capillary. During phase III, the sphincter constricted significantly more than the PA and the bulb. Datasets were analyzed via LME models followed by Tukey post hoc tests for pairwise comparisons (Phase II data was log-transformed to ensure homoskedasticity). **e**, Boxplot summary of the estimated flow resistances at vessel segments during baseline and phase III of CSD. Paired Wilcoxon signed rank tests were used to establish difference (*p* < 0.05) before and during CSD phase III.

We also examined the vulnerability of the precapillary sphincter to global ischemia induced by cardiac arrest (Fig. 6). Cardiac arrest caused an immediate loss of blood pressure and an initial 26±7% mean drop in lumen diameter within the first 2 minutes (Fig. 6b,c, n = 7), which was most prominent at the bulb. Over the subsequent 30 minutes we observed vasoconstriction of the cerebral microvessels that occurred with important differences in time delay. The sphincter remained relatively unchanged for the first ~14min, whereas the PA and 1^st^ order capillary showed a steady reduction in diameter. After 14-20 minutes, we observed an accelerated constriction spreading from the 1^st^ order capillaries towards the PA and along the PA towards the brain surface (Supplementary video 7). The precapillary sphincter collapsed at a rate of 0.23±0.03 μm min^−1^ (Fig 6c, phase II). The collapse of the sphincter was complete after ~25min and caused an extreme increase in flow resistance (Fig 6d, phase III) that essentially occluded entry of RBCs into the capillary networks. Concurrent with the collapse of the vessel lumen, we observed a swelling of astrocytic end-feet and vasculature-associated astrocyte soma (Supplementary video 7).

**Figure 6.**
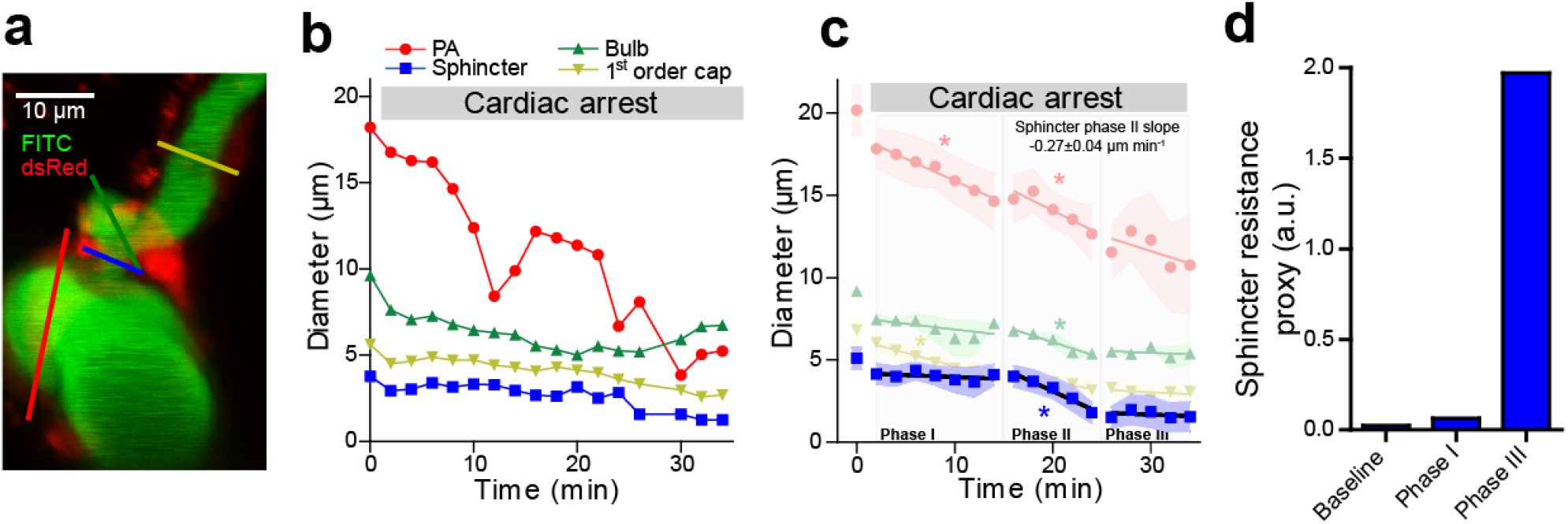
The integrity of precapillary sphincters represents a vulnerable site to prolonged ischemia. Cardiac arrest and global ischemia was elicited by intravenous injection of pentobarbital (50 μl of a mix of 200 mg/mL pentobarbital and 20mg/mL lidocaine) during imaging of the precapillary sphincter. **a**, Representative maximal intensity projection of a FITC-dextran loaded NG2-dsRed mouse. Color coded lines mark the ROIs for repeated diameters measures before and after cardiac arrest. **b**, Representative time series of diameter changes at vessel segments during cardiac arrest. The initial diameter decrease is due to the loss of blood pressure. **c**, Summary time series of diameter changes after cardiac arrest. Three phases were constructed based on the dynamics of the sphincter. In phase I, the sphincter and bulb diameters held steady, while the PA and 1^st^ order capillary diameter gradually declined. After ~15 min (phase II), both the sphincter and bulb starts to collapse. After ~25min (phase III), the sphincter is collapsed and only the PA shows continuing decline. * indicates a slope of the linear regression significantly different from zero, n = 3-7 mice. **d**, Estimates of changes in flow resistance within the sphincter using Poiseuille’s law at baseline, in phase I, and in phase III.

## Discussion

The organization of the cortical vasculature simultaneously accommodates enough pressure for proper perfusion of each cortical layer while preventing the pressure head from inducing tissue damage. Here, we show that precapillary sphincters exist and that they represent active bottlenecks that are strategically located at the upper part of the cortex preferentially in larger PAs that branch into large capillaries. This localization contributes to equalization of perfusion along the length of the PA. In addition, the pressure drop across the narrow sphincter (~2-5 μm) protects downstream capillaries and brain tissue both under baseline conditions and during functional stimulation (Fig. 2–4). However, the high sensitivity of flow resistance to constriction becomes precarious in pathological conditions that promote general constriction (Fig. 5–6).

### The precapillary sphincter equalizes perfusion along the PA and protects downstream tissue against adverse pressure spikes

Precapillary sphincters abound at proximal branches of relatively large PAs descending to relatively large 1^st^ order capillaries, primarily in upper cortical layers (Fig. 2), i.e. in the cerebral microvessels that withstand the high arterial pressure. The sphincters have high flow resistances and reduce the transmural pressure in downstream capillaries, protecting the endothelium from disruption, and contributes to equalize perfusion to all capillary networks along the entire length of the PA. The sphincter location is consistent with the assumption that the blood pressure drop at 1^st^ order capillaries is largest at superficial layers and decrease over the depth of the cortex^25^. Precapillary sphincters represent a bottleneck that may protect the capillary networks from mechanical impact while at the same time preserving the perfusion pressure in the penetrating arteriole along the entire cortical depth. Our data are consistent with studies from other groups which indicate that regulation of capillary blood flow can occur independently from arteriolar flow control^2,26,27^.

### The distention at the bulb decelerates RBCs, possibly providing a mechanism of RBC alignment in capillaries

The bulb, i.e. the distention that usually appeared immediately downstream from the precapillary sphincter (Fig. 1), had areas lacking NG2-expression (Extended data Fig. 4 and Supplementary video 2) which may indicate sparse pericyte coverage. Similar to previous observations^28^, we consistently observed short and thick endothelial nuclei at the bulb (Extended data Fig. 4 and Supplementary video 2). The bulb remained less vasoactive compared to the precapillary sphincter and 1^st^ order capillary, consistent with less contractile cell coverage (Fig. 1a, Extended data Fig. 4 and Supplementary video 2). The large cross-sectional area of the bulb caused deceleration and deformation of RBCs from a bullet to parachute form^29^ (Supplementary video 5) followed by realignment as RBCs entered the capillary network.

### The precapillary sphincter is a highly active flow regulator but remains limited by passive structural elements

Our initial analysis of the sphincter complexes suggested a dual role: first, in distributing pressure and perfusion along the PA and second by protecting brain tissue and downstream capillaries against adverse pressures and hemorrhage. In principle, these functions might arise from both active contractile elements and passive structural elements. α-SMA protein is key for contractile function and is widely expressed in vascular smooth muscle and routinely found in pericytes of 1^st^ order capillaries within cortex^14,15,30^. In accord with previous data^31^, we observed α-SMA along both the PA and 1^st^-4^th^ order capillaries and importantly, within the pericyte constituting the sphincter (Fig. 3a). The presence of α-SMA explains the capacity for active vasomotor responses at the sphincter (Fig. 3c-i). The integrity and morphology of the sphincter remained preserved after local administration of papaverine despite a significantly larger dilation for the sphincter as compared to the bulb and 1^st^ order capillary (Fig. 4c)^20^. We report elastin^20^ (Fig. 4e, Alexa 633 hydrazide) and filamentous collagen α1 type 1 (Fig. 4f, COL1A1) expression that may support the structural integrity of the sphincter during rises in blood pressure (Fig. 4d,e). The high capacity for diameter variations at the sphincter during functional stimulation suggests that cortical flow control resides both in capillaries and at arteriole branchpoints and may reconcile some of the controversies regarding the dynamic regulation of cerebrovascular resistance as described previously^2,15,30,32^. Sphincters are located at proximal PA branches and their occurrence suggests that regulation of cerebral blood is distributed between arterioles and capillaries depending on the local angioarchitecture (Fig. 2). It follows, that the distribution of flow resistance may shift dynamically with changes in perfusion demand^33–35^ (Fig. 3). Furthermore, during functional sphincter dilation only one RBC at a time passed into the bulb and 1^st^ order capillary, which suggests that sphincters contribute to plasma skimming^36,37^, and thereby contribute to redistribution of RBCs and (in consequence) of hematocrit within the local vascular network.

### The precapillary sphincter is vulnerable to pathological conditions of general constriction

Cortical spreading depression (CSD) is a slow depolarizing wave along the cortex that is involved in migraine, traumatic brain injury, and stroke^38^. CSD evokes an initial vasoconstriction (phase I), immediately followed by a transient hyperemic response (phase II), which is superseded by a long lasting vasoconstriction of arterioles and capillaries (phase III) during which the neurovascular coupling is impaired^4,39^. During a CSD, the sphincter displayed pronounced diameter changes (Fig. 5) and constricted persistently during the long period of low blood flow after CSD (Supplementary video 6). Persistent sphincter constriction reduced both the RBC flow rate and the hematocrit of the capillary bed. The long lasting oligemia previously described in CSD could arise from the high resistance observed at precapillary sphincters^4^, and further pharmacological research on this structure could improve help the outcome of CSD in the ischemic brain or in migraine patient. We also examined the reaction of sphincters to cardiac arrest (Fig. 6). We have previously shown that cerebral pericytes during simulated global ischemia immediately begins to constrict and later starts dying in rigor after ~15 min in rat cortical brain slices^2^ (all pericytes lost after ~40min). During *in vivo* imaging of cardiac arrest, we observed a steady vasoconstriction of the precapillary sphincters around 16 min after onset of cardiac arrest (Fig. 6c) and collapse at ~25 min. The other vessel segments displayed linear diameter reductions but no collapse. During global ischemia, we also observed swelling of astrocyte end-feet and soma, which are known to compress microvessels^40^, probably adding to the sphincter collapse (Supplementary video 7).

## Conclusions

Precapillary sphincters represent important anatomical sites of blood flow regulation due to their strategic placement at branchpoints of proximal PA’s, where they control perfusion along the PA. We show that while maximal dilation of the sphincter pericyte is structurally limited, it displays a high capacity of vasomotor activity around a baseline diameter of 3-4 μm, where flow resistance is most sensitive to diameter changes. Therefore, precapillary sphincters represent a mechanism to equalize pressure and RBC flux between the capillary networks that branch off from the upper, middle, and lower parts of the PA. At the same time, sphincters protect downstream capillaries and brain tissue against adverse blood pressure spikes. The dual function is crucial for proper perfusion of cortical vessels. During pathology, sphincter constriction severely limits perfusion of downstream capillaries in CSD and prolonged ischemia. Prevention of sphincter constriction may be of therapeutic importance in migraine and cerebral ischemia.

## Methods

### Animal Handling

Animal procedures were approved by The Danish National Ethics committee according to the guidelines set forth in the European Council’s Convention for the Protection of Vertebrate Animals Used for Experimental and Other Scientific Purposes. 32 male or female NG2-dsRed mice (Tg(Cspg4-DsRed.T1)1Akik/J; Jackson Laboratory; 19 to 60 weeks old) and 27 male or female wild-type mice (C57bl/6j; Janvier-labs, France;16 to 32 week) were used. The NG2-DsRed mice were used in the studies of whisker pad stimulation, cardiac arrest, thinned skull and local ejection of papaverine. The rest of the studies were performed in wild-type mice.

### Surgical Procedures

Anesthesia was induced with bolus injections of xylazine (10mg/kg, intraperitoneally (i.p.)) and ketamine (60 mg/kg, i.p.) and maintained during surgery with supplemental doses of ketamine (30mg/kg/20 min, i.p.). Mechanical ventilation (Harvard Apparatus, Minivent type 845) was controlled through a cannulation of the trachea. One catheter was inserted in the left femoral artery to monitor blood pressure and to collect blood samples. Another cathether was inserted in the femoral vein to administer chemical compounds. The content of blood gasses in arterial blood samples (50 μl) was analyzed by an ABL700 (Radiometer, Copenhagen; pO2, normal range: 95–110 mmHg; pCO2, normal range: 35–40 mmHg; pH, normal range: 7.35–7.45). To maintain physiological conditions, both respiration and mixed air supply was adjusted according to the blood gas analysis or occasionally according to continuously monitored end-expiratory CO2 (Harvard Apparatus, Capnograph 340), blood oxygen saturation (Kent Scientific, MouseStat pulsoximeter). A craniotomy (diameter ~3 mm. Center coordinates: 3 mm right of and 0.5 mm behind bregma) was drilled above the right somatosensory barrel cortex. We switched anesthesia to α-chloralose (33% w/vol; 0.01 mL/10 g/h) upon surgery completion. In the end of the experiments, mice were euthanized by intravenous injection of pentobarbital followed by cervical dislocation.

To ensure that precapillary sphincters were not a result of the craniotomy, we made thinned skull preparations over the barrel cortex, at the point of the surgical procedure where we would otherwise have made a craniotomy. By following the protocol by Shih et al^41^, we thinned the skull to around 40 μm thickness, polished with tin oxide powder and covered the window with agarose and a coverslip.

### Chronic cranial window implantation

A chronic cranial window was installed approximately three weeks prior to imaging in mice of a C57Bl/6 background. The surgical procedure is adapted from Goldey et al.^42^. A small craniotomy was performed over the left barrel cortex under isoflurane anaesthesia and a custom-made reinforced cover glass consisting of three 3 mm coverslips glued on top of each other and onto a 5 mm coverslip was installed. A custom-made head bar was attached to the right side of the skull allowing for head immobilization during imaging sessions. In the five days following implantation the animal was closely monitored and given pain and infection treatment as described in Goldey et al.^42^. When the animal had recovered after surgery, training for imaging experiments could commence. The animal was familiarized with the experimenter through gentle handling. After several handling sessions and when the animal was comfortable with the experimenter, it was slowly accustomed to head immobilization. The animal was given treats in form of sweetened condensed milk during the training process. When the animal had been habituated with the head immobilization for periods of about an hour of length, they are ready for imaging experiments.

### Electrical stimulation in whisker pad

The mouse sensory barrel cortex was activated by whisker pad stimulation. The contralateral ramus infraorbitalis of the trigeminal nerve was electrically stimulated using a set of custom-made bipolar electrodes inserted percutaneously. The cathode was positioned relative to the hiatus infraorbitalis (IO), and the anode was inserted into the masticatory muscles. Thalamocortical IO stimulation was performed at an intensity of 1.5 mA (ISO-flex; A.M.P.I.) for 1 ms in trains of 20 s at 2 Hz.

### Pressure ejection of papaverine via glass micro-pipette

Borosilicate glass micro-pipettes were produced by a pipette puller (P-97, Sutter Instrument) with a resistance of 2.5 ~ 3.0 MΩ. The pipette was loaded with a mixture of 10 μM Alexa 594 and 10 mM papaverine in order to visualize the pipette tip in both epi-fluorescent camera and two-photon microscope. Guided by the two-photon microscopy and operated by a micromanipulator, the pipette was carefully inserted into the cortex to minimize tissue damage and avoid vessel bleeding. The distance between the pipette tip and vasculature was 30-50 μm. Papaverine was locally ejected for ~1 s by 3 times using an air pressure of <15 psi in the pipette (PV830 Pneumatic PicoPump, World Precision Instruments). A red cloud (Alexa 594) ejected from the pipette tip was visually observed to cover the local vascular region simutaneously, and the background returned to normal approximately 1 minute after puffing^3^. Papaverine was pre-conditioned for 5 minutes before imaging the same vasculature again.

### Cortical Spreading Depression

In a subset of experiments, cortical spreading depolarization (CSD) was triggered 2 mm away from the recording site using pressure injection of 0.5 M potassium acetate (KAc) into the cortex (estimated volume ~0.5 μl). Apart from triggering CSD, KAc injection did not cause a brain lesion (bleeding or tissue damage), and to ensure that no spreading depolarization was triggered during surgical preparation, our technique for making craniotomies has previously been validated by measuring cerebral blood flow while drilling the craniotomy. Using this technic, we could show that no spreading depolarization was elicited^39^. Moreover, prior to our first spreading depolarization we measure whisker responses, and mice that did not show normal vessel diameter responses to stimulation were discarded from the dataset, ensuring that no spreading depolarization were triggered before recordings started.

### Two-photon imaging

Images and videos were obtained using two sets of laser-scanning two-photon microscopes. Experiments of prevalence quantification, RBC velocity, evoked Ca^2+^ responses, CSD, and cardiac arrest were performed using a commercial two-photon microscope (FluoView FVMPE-RS, Olympus) equipped with a 25 x1.05 NA-water-immersion objective (Olympus) and a Mai Tai HP Ti:Sapphire laser (Millennia Pro, Spectra Physics). Experiments of whisker pad stimulation and papaverine ejection were performed using a second commercial two-photon microscope (Femto3D-RC, Femtonics Ltd.) with a 25 × 1.0 NA water-immersion objective with piezo motor and a Ti:Sapphire laser, Mai Tai HP Deep See, Spectra-Physics. The excitation wavelength was set to 900 nm. The emitted light was filtered to collect red (590–650nm) and green (510–560nm) light from dsRed (pericytes) or SR101 (astrocytes) and FITC-dextran (vessel lumen) or OGB (relative Ca^2+^ changes), respectively.

The prevalence of precapillary sphincters and bulbs were studied by acquiring image stacks using our Olympus two-photon microscope (Fluoview), tracking each penetrating arteriole from pial to >650 μm in depth in a frame-scan mode at around 1 frame per second with pixel resolution of 512×512 at 1000 nm excitation wavelength. Measurements of RBC velocity were performed in resonance bi-directional line-scan mode with a scan rate of 15,873 Hz (0.063 ms per line) and pixel resolution of 512 pixels per line. Evoked Ca^2+^ responses and CSD was imaged in one vessel branching from the penetrating arteriole to a first order capillary, including the neck and bulb structure in a single plane. Evoked Ca^2+^ responses and CSD’s was imaged in one vessel branching from the penetrating arteriole to a first order capillary, including the neck and bulb structure in a single plane. The excitation wavelength was set to 900, the frame resolution was 0. 450 μm/pixel with a 320 × 240 pixels frame and images were taken at a speed of 2.40 frames per second for evoked Ca^2+^ responses. The excitation wavelength was set to 920 nm, the frame resolution was 0.255 μm/pixel with a 512 × 384 pixels frame and images were taken at a speed of 0.81 frames per second for CSD.

In the experiments using Femtonics microscope, we recorded the whole volume including the vessel segments of interest by fast repetitive hyperstack imaging (4D imaging) – continuous cycles of image stacks along the z-axis. This is to compensate focus drift and study vasculatures spanning in a certain z-axis range. Each image stack was acquired within 1 second that comprised 9-10 planes with a plane distance of 3 – 4 μm. This covered the whole z-axis range of the investigated blood vessels. The pixel sizes in the x-y plane were 0.2 – 0.3 μm.

### Two-photon imaging analysis

Data were analyzed in ImageJ or MATLAB using custom-built software. In the study of prevalence of precapillary sphincters and cardiac arrest, multiple ROIs were manually placed across vessel lumen in ImageJ, measuring vessel diameters. In the study of whisker pad stimulation induced evoked Ca^2+^ responses, ROIs were detected using a modification of the pixel-of-interest-based analysis method^43^. ROIs were positioned around astrocytic end-feet and labeled according to their location on PA, sphincter or 1^st^ order capillary. Further ROIs were positioned around neuronal somas and in neuropil. Astrocytic or neuronal structures were recognized based on SR101/OGB staining, cell morphology, and relation to blood vessels^44^. For each frame, we selected pixels showing intensities of 1.5 standard deviations (SD) above the mean intensity of the ROI. The intensities of these pixels were averaged and then normalized to a 15-s baseline period just before stimulation onset, creating a time trace of ΔF/F0 for every ROI. These time traces were smoothed with a 3-sec moving average to avoid outlier values. A Ca^2+^ transient was defined as an intensity increase of ≥5% and of ≥2 SD from baseline, having a duration of ≥2.5 s. Recordings were divided into 20-s time bins (baseline, dilation, undershoot). Responsivity was defined as the fraction of ROIs with Ca^2+^ transients within time bins per mouse. Recordings lacking evoked Ca^2+^ responses in neuropil were excluded from analysis. In the study of CSD, rectangular region of interests (ROIs) with width of 2 or 4 μm were drawn perpendicular to the surface of the vessel at the defined locations. An active contour algorithm (Chan-Vese segmentation) was used to calculate the vessel diameter change in these ROIs. The diameter change over time was detected for each ROI. For the 4D imaging performed in whisker pad stimulation and papaverine ejection, each image stack was flattened onto one image by maximal intensity projection, which converts the data to the same formats of CSD. Similar diameter analysis methods were used. Values from each ROI type were averaged per mouse. Arteriole bifurcations leading to two equally sized arterioles and 1^st^ order capillaries bifurcating under 10 μm from the arteriole branchpoint were not included in the analysis. 3D renderings for supplementary videos were done with Amira software (ThermoFischer scientific).

### Immunohistochemistry

Adult NG2-ds-Red mice were transcardially perfused with 4% paraformaldehyde (PFA) and their brains were extracted and cryoprotected in 30% sucrose, rapidly frozen in cold isopenthane (−30°C) and sectioned using cryostat at 25 and 50 μm thickness. Sections were rinsed (3 × 5 min) in 0.1 M PBS and antigen retrival was performed (for Collagen-I staining) using hot citrate buffer (90°C, pH 6.0) for 20 minutes. 50 μm sections were permeabilized and blocked in 0.5% Triton-X 100 in 1 X PBS (pH 7.2) and 1% BSA overnight at 4°C, whereas 25 μm sections were permeabilized in 0.5% Triton-X 100 in 1 X PBS for 30 minutes and blocked in 5% NGS, 5% BSA and 0.5% Triton-X 100 in 1 X PBS for 1 hour at RT. Sections were incubated in primary antibodies for two nights at 4°C in blocking buffer containing 1-5% BSA, 5% NGS in 0.25-0.5% Triton-X 100 in 1 X PBS. The following primary antibodies were used: mouse ACTA2-FITC (1:200; Sigma; F3777) and rabbit anti-Collagen I (1:50; ab34710). Elastin was labeled using an artery-specific red dye; Alexa Fluor 633 (A30634, ThermoFisher Scientific) at 1:300 dilution from 2mM stock. Alexa Fluor 633 was added to the brain sections for 10 minutes and rinsed. Thereafter, the sections were washed (3 × 5 min) in 0.1 M PBS and incubated with goat anti-rabbit Alexa488 (1:500; Sigma-Aldrich) secondary antibody (to label collagen-I) for 1 hour at RT. After incubation with secondary antibody sections were rinsed (3 × 5 min) in 1X PBS, incubated in Hoechst (1: 6000) for 7 minutes, rinsed again (3 × 5 min) in 1X PBS and mounted using SlowFade™ Diamond Antifade Mountant (Invitrogen; S36963). Fluorescence images were acquired with a confocal laser scanning microscope (LSM 700 or 710) equipped with Zen software at 20x/0.8 NA and 63x/1.40 NA oil DIC M27 objectives, 1X (0.170 μm/pixel) and 4X (0.021 μm/pixel) digital zoom respectively. Care was taken to ensure similar fluorescence across images.

### Statistical analysis

Datasets are presented as mean ± s.e.m, standard box plots, or in case of log-transformed data as back-transformed means ± 95% confidence intervals. Normality of data was assessed using Shapiro-Wilk and graphical tests. For normal datasets, Linear Mixed Effects (LME) model analyses were performed. LME was chosen to take proper advantage of multiple measurements of parameters and/or of multiple time points in the same animal. Vessel segments (PA, sphincter, bulb, and 1^st^ order capillary) were included as the fixed effect, whereas the particular mouse and vessel branch were included as random effects as needed. Heteroscedastic datasets were log-transformed to conform to analysis as indicated. Significant difference (p-value < 0.05) was obtained by likelihood ratio tests of the LME model with the fixed effect in question against a model without the fixed effect. Tukeys post-hoc test was used for pairwise comparisons between elements in the fixed effect group. For non-normal data, non-parametric Wilcoxon signed-rank tests were used for paired samples, while the Kruskal-Wallis test was used for multiple independent groups; for pairwise comparisons, the Wilcoxon rank-sum test with Holm’s p-value adjustment method was used. Finally, linear regression was used to assess the relationships were fitted to datasets. All statistical analysis was conducted using R (version 3.4.4; packages *lme4*^45^ and dplyr) and Prism version 5.

## Supporting information

Video of a thinned skull in vivo experiment

Z-stack and volume rendering of a smooth muscle actin antibody and DAPI staining at a precapillary sphincter of an NG2-dsRed mouse.

Hyperstack projection of a precapillary sphincter dilation after whisker pad stimulation.

Video of RBCs getting stuck temporarily at the precapillary sphincter

Resonance frame-scan video of RBCs passing through a precapillary sphincter and taking up the parachute form.

Video of the three phases of the CSD

Hyperstack recording of precapillary sphincters in an NG2-dsRed or WT mouse (+SR101) mouse after cardiac arrest.

## Acknowledgements

We would like to acknowledge our animal technician Micael Lønstrup for his help with animal experiments. Nikolay Kutuzov and Dr. Krzysztof Kucharz for scientific discussions. Thanks to Kirsten Thomsen for advices regarding statistical analysis. A special thanks to Prof. Anna Devor for hosting and supervising Aske Graakjær Krogsgaard Jensen during his visit in San Diego working with awake mice. Thanks to the core facility for integrated microscopy (CFIM) at our institute for their service.

