## Extended data for "Precapillary sphincters control cerebral blood flow"

#### Extended data table 1 – Whisker stimulated vasodilation

| | PA | Sphincter | Bulb | 1st order capillary |
|---|---|---|---|---|
| **Baseline diameter (μm)** | 18.43±1.57 | 4.55±0.37 | 7.90±0.64 | 6.48±0.63 |
| **Max vasodilation (%)** | 10.97±2.27 | 33.75±4.08 | 12.92±3.98 | 15.53±2.27 |
| **Max undershoot (%)** | -2.08±0.85 | -12.40±2.10 | -8.48±3.22 | -6.71±2.45 |

Data shown as mean ± s.e.m. See Fig. 3 for statistics.

#### Extended data calculation 1 – RBC flux calculation

The RBC count of an adult mouse blood[1] is around $9 \cdot 10^6$ cells/mm$^3$ and RBC velocity ($V_{RBC}$) through the sphincter (radius, r ≈ 1.75 µm) is measured to be 8.7 mm/s at baseline. Disregarding the propensity of lower hematocrit in the microcirculation due to plasma skimming, a rough calculation gives

$$RBCflux = \pi \cdot r^2 \cdot V_{RBC} \cdot RBC_{count} = RBC/s$$

This value is somewhat higher than the 543 cells/s measured during baseline, which probably reflects the lower hematocrit of cortical blood compared to the reference count. Note, however, that the calculated value is highly sensitive to the values used but does show that a high number of RBCs pass through a sphincter every second.

#### Extended data figure 1 – Illustration of *in vivo* setup:

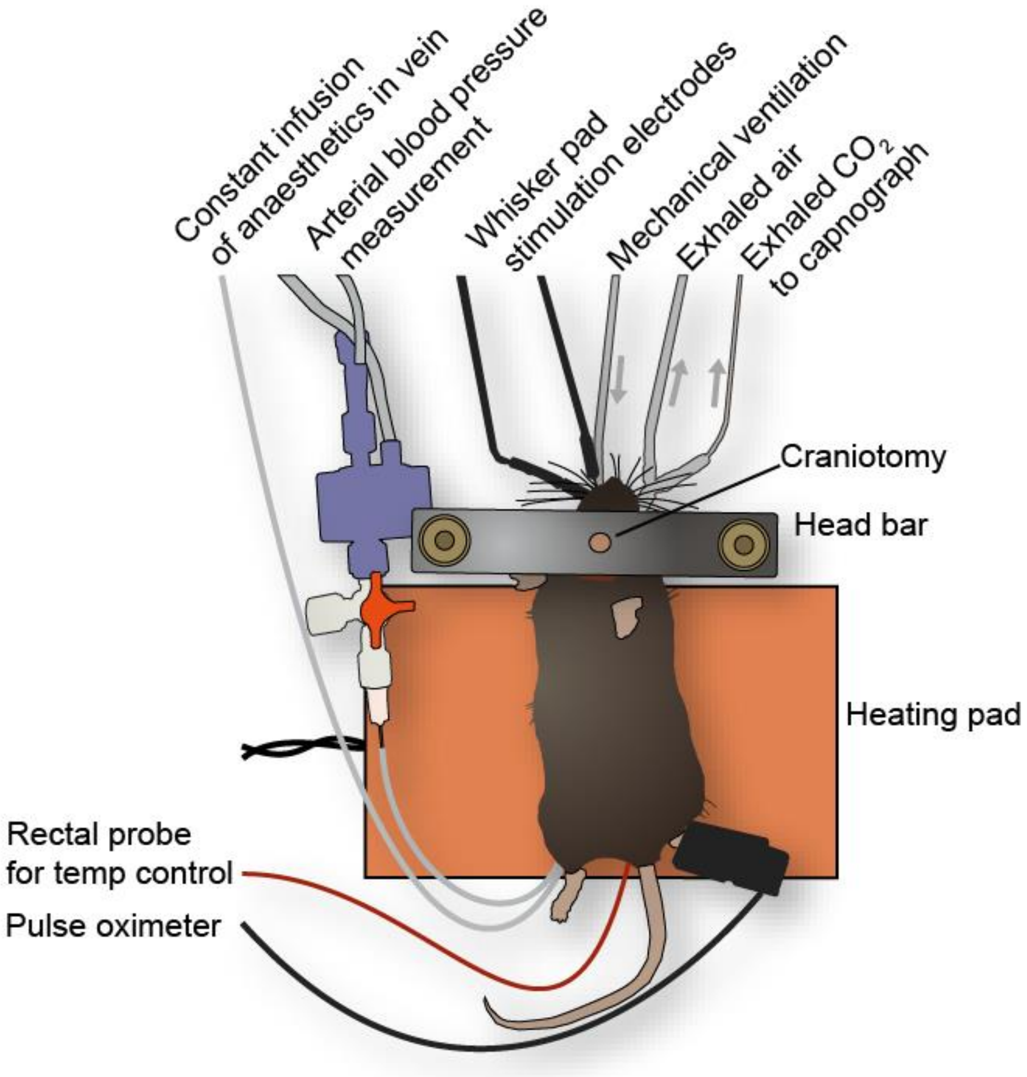


*Extended data Figure 1 | Illustration of the in vivo setup. The mouse was mechanically ventilated, anaesthetized with α-chloralose infusion in the left femoral vein (only after surgery) and kept warm with a heating pad with a rectal probe as reference. To monitor the physiological state of the animal we measured exhaled $CO_2$ with a capnograph, measured the left femoral artery blood pressure with a blood pressure monitor and measured the oxygen saturation with a pulse oximeter. We also took blood samples to check the blood gasses and corrected ventilation and/or oxygen saturation of the mixed air to keep the animal in a physiological state. The mouse skull was glued to a head bar and a 3-mm craniotomy was drilled over the right barrel cortex. Stimulation electrodes were inserted into the contralateral ramus infraorbitalis of the trigeminal nerve.*

### Extended data figure 2 – Precapillary sphincters in thinned skull preparation:

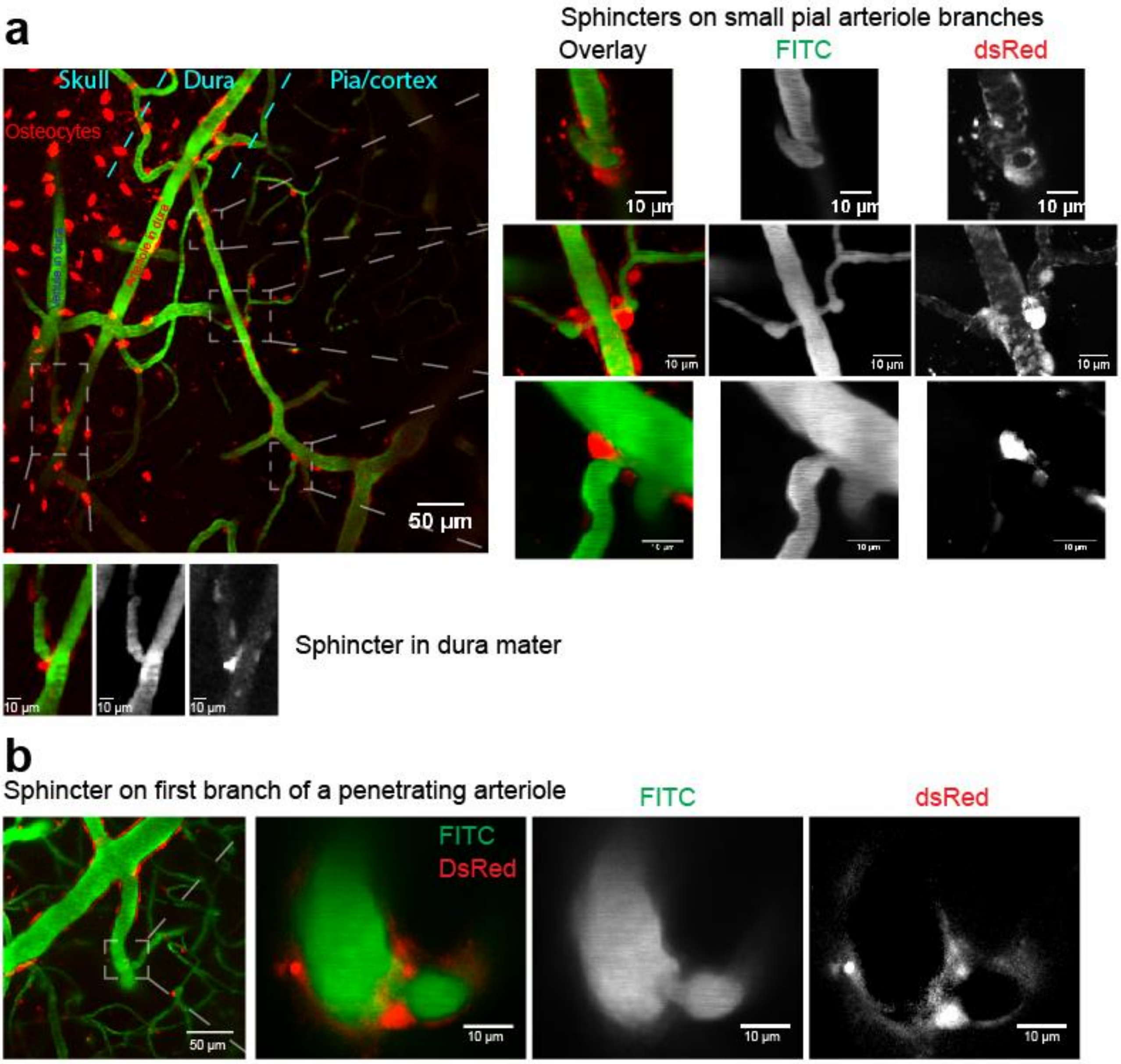


*Extended data Figure 2 | Two photon imaging through thinned skulls of dsRed mice injected with FITC.* ***a***, ***Left panel****: Example of a maximum intensity projection of a thinned skull experiment, where the dsRed positive osteocytes of the skull are seen in the left side (the image is taken on an angle, see Supplementary video 1), fading to the right into the blood vessels in the dura and then to the far right fading into a pial arteriole and cortical capillaries.* ***Right panels****: Local projections of three precapillary sphincters on surface vessels (see also Supplementary video 1 and 4).* ***Lower panels****: Example of a precapillary sphincter on a dural arteriole branch.* ***b,*** *Another example of a thinned skull experiment, showing a maximim intensity projection of a penetrating arteriole, and to the right a local projection of the precapillary sphincter on the first branch.*

#### Extended data figure 3 – Precapillary sphincters in an awake mouse:

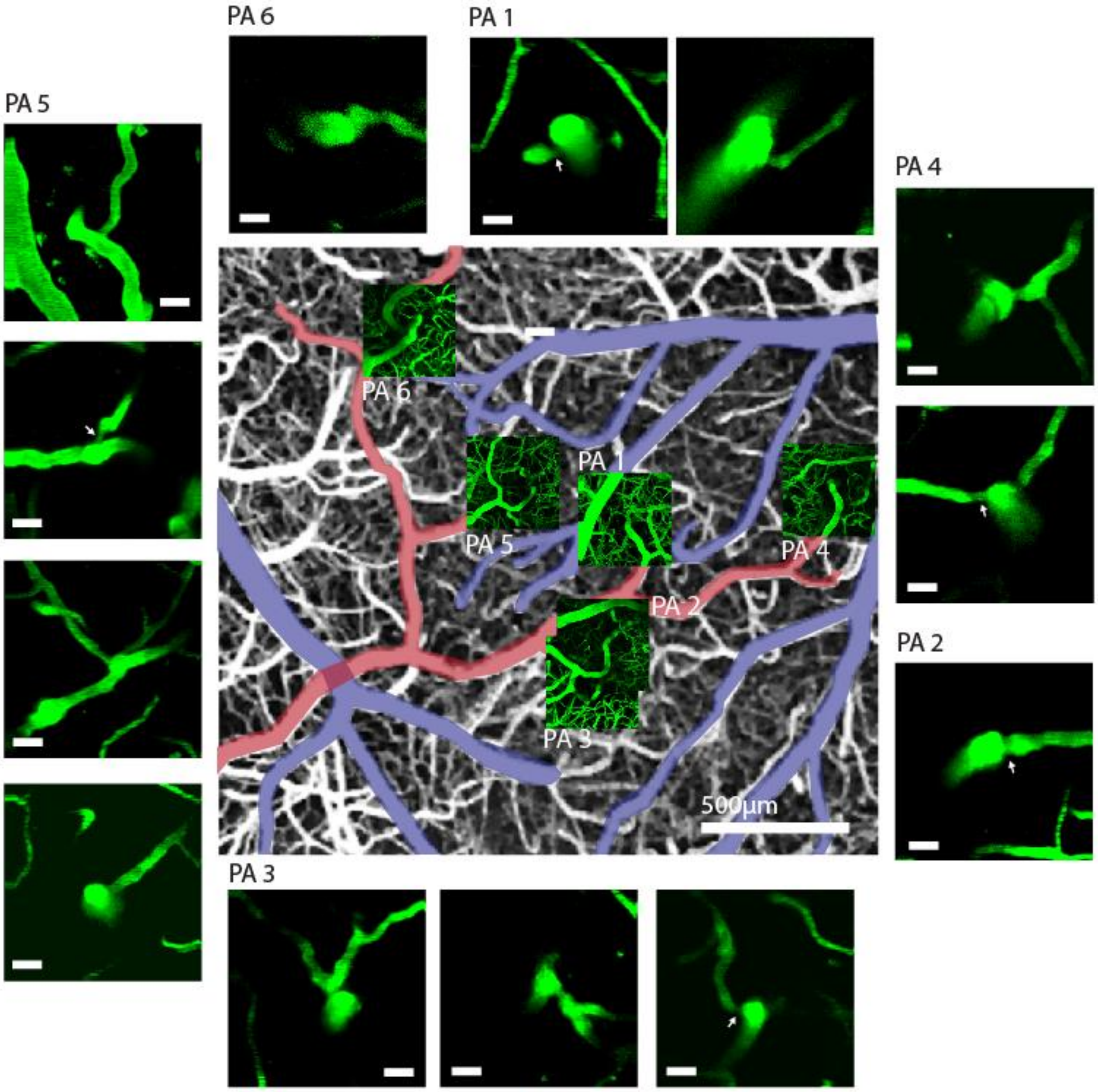


*Extended data Figure 3 | **Center**: Part of whisker barrel cortex in a mouse of a C57bl6/j background. Vessels have been loaded with FITC-dextran. Veins have been colored blue and arteries red. Maximum intensity projection of the uppermost 300 μm of cortex surrounding 6 penetrating arterioles is shown in location. **Surrounding**: Maximum intensity projections of local z-stacks of branching points on the six penetrating arterioles are shown. The first row contains the uppermost branching points. The second row the second branching point and so on. The scalebars are 20 μm. White arrows mark precapillary sphincters, precapillary sphincters on bifurcating PA branches are not marked.*

Extended data figure 4 – Smooth muscle actin stains up to 4$^{th}$ order capillaries:

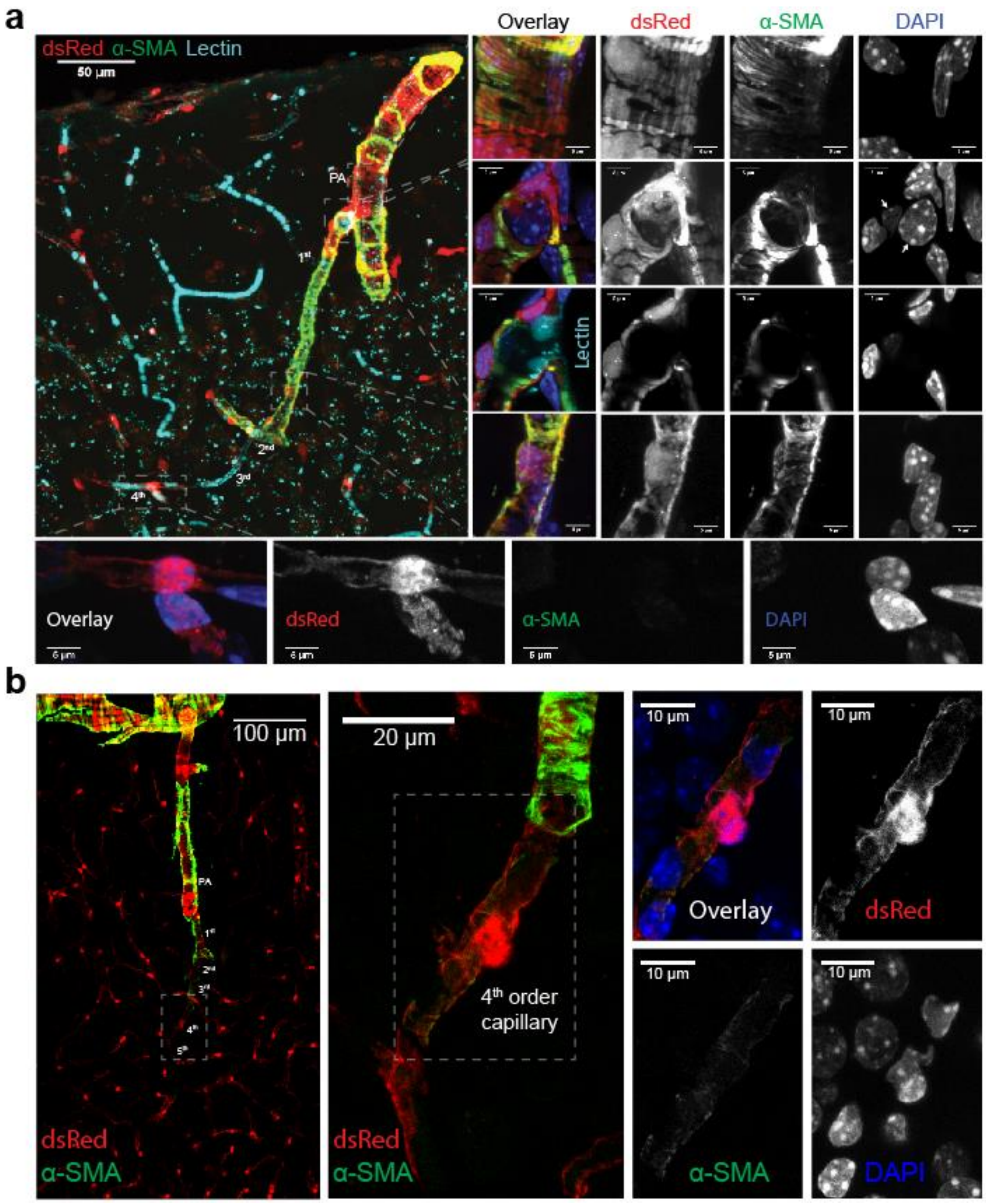


*Extended data Figure 4 | Immunostaining for α-smooth muscle actin in coronal slices of NG2-dsRed mice.* ***a****,* ***Top left****: Maximum projection image of a penetrating arteriole in a mouse where we injected tomato-lectin (cyan) 20 min before PFA fixation. Notice that the tomato-lectin also stains residual RBC's in the capillaries.* ***Top right****: local projections of the PA showing filamentous for α-SMA in smooth muscle/pericyte hybrids, below the precapillary sphincter showing α-SMA expression in the sphincter pericyte, an uncovered endothelial nucleus at the bulb and α-SMA expression in the 1$^{st}$ order pericyte. Below, a narrow projection (cross section) of the of the precapillary sphincter including the lectin stain, showing the narrow lumen of the precapillary sphincter and the large volume of the bulb. Below a maximum projection of the far end of the 1$^{st}$ order capillary.* ***Lower panel****: Maximum intensity projection of a 4$^{th}$ order capillary showing no α-SMA expression.* ***b****, Another example of α-SMA immunostaining of a penetrating arteriole in a dsRed mouse, showing α-SMA expression up until the 4$^{th}$ order capillary. To the right are magnifications of the 4$^{th}$ order capillary showing weak but existing α-SMA expression of an ensheathing pericyte.*

Extended data figure 5 – Resistance at the precapillary sphincter:

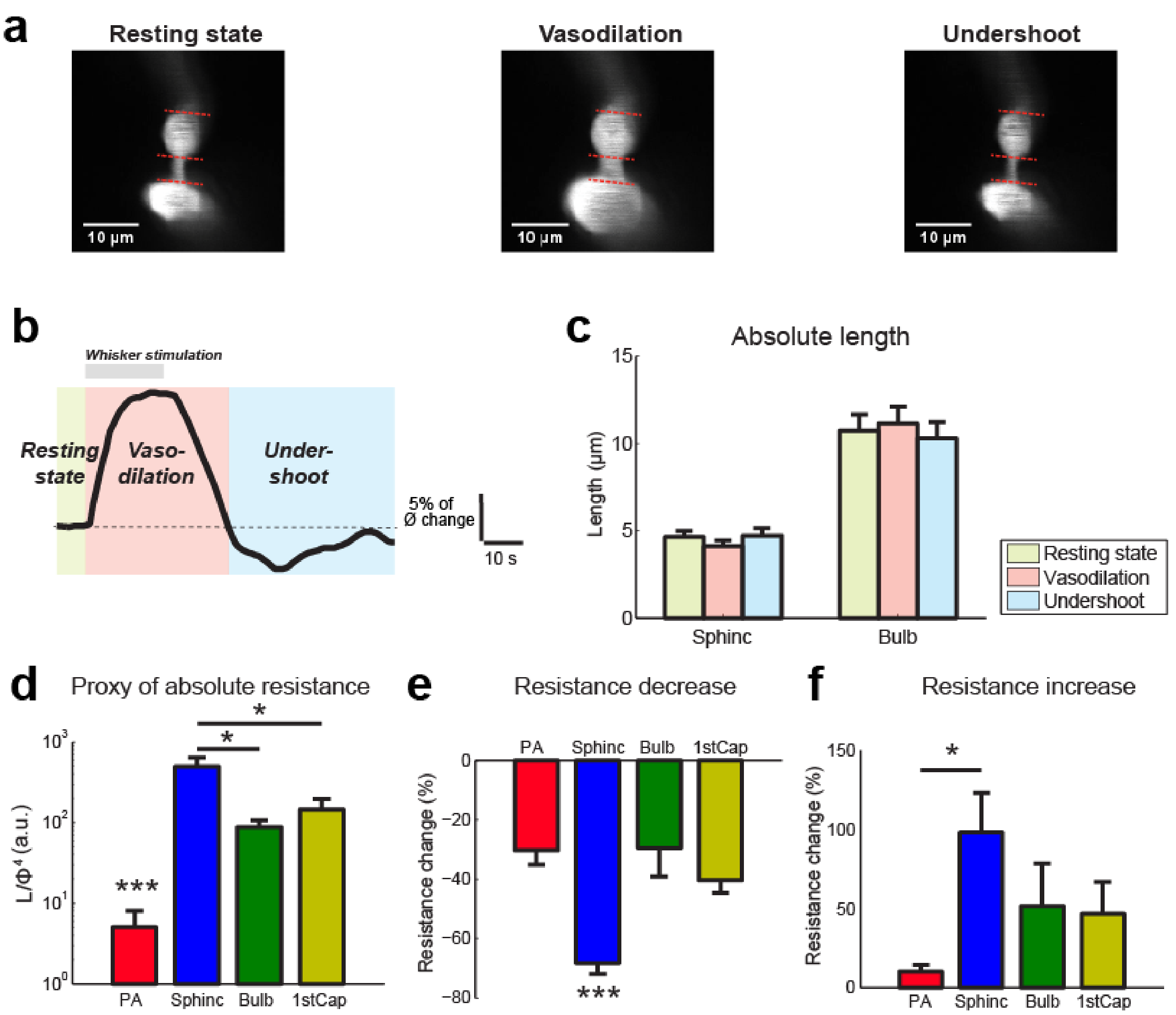


*Extended data Figure 5 | Precapillary sphincter length decreases during stimulation, augmenting the decrease of resistance.* ***a****, Two-photon images of a precapillary sphincter and bulb branching from a PA (lower part), red lines indicate segments edges of sphincter and bulb at baseline.* ***b****, Illustration of the three phases of vasodilation.* ***c****, Absolute lengths of sphincter and bulb in the three phases of vasodilation.* ***d****, Poiseuille's law derived proxy of resistance including the length of the segment at baseline (the length of the PA and 1st order capillary is set to the length of the sphincter).* ***e****, Resistance decrease during vasodilation after inclusion of the length change in the proxy of resistance.* ***f****, Resistance increase during undershoot after inclusion of the length change in the proxy of resistance.*

Extended data figure 6 – Evoked **$Ca^{2+}$ responses:**

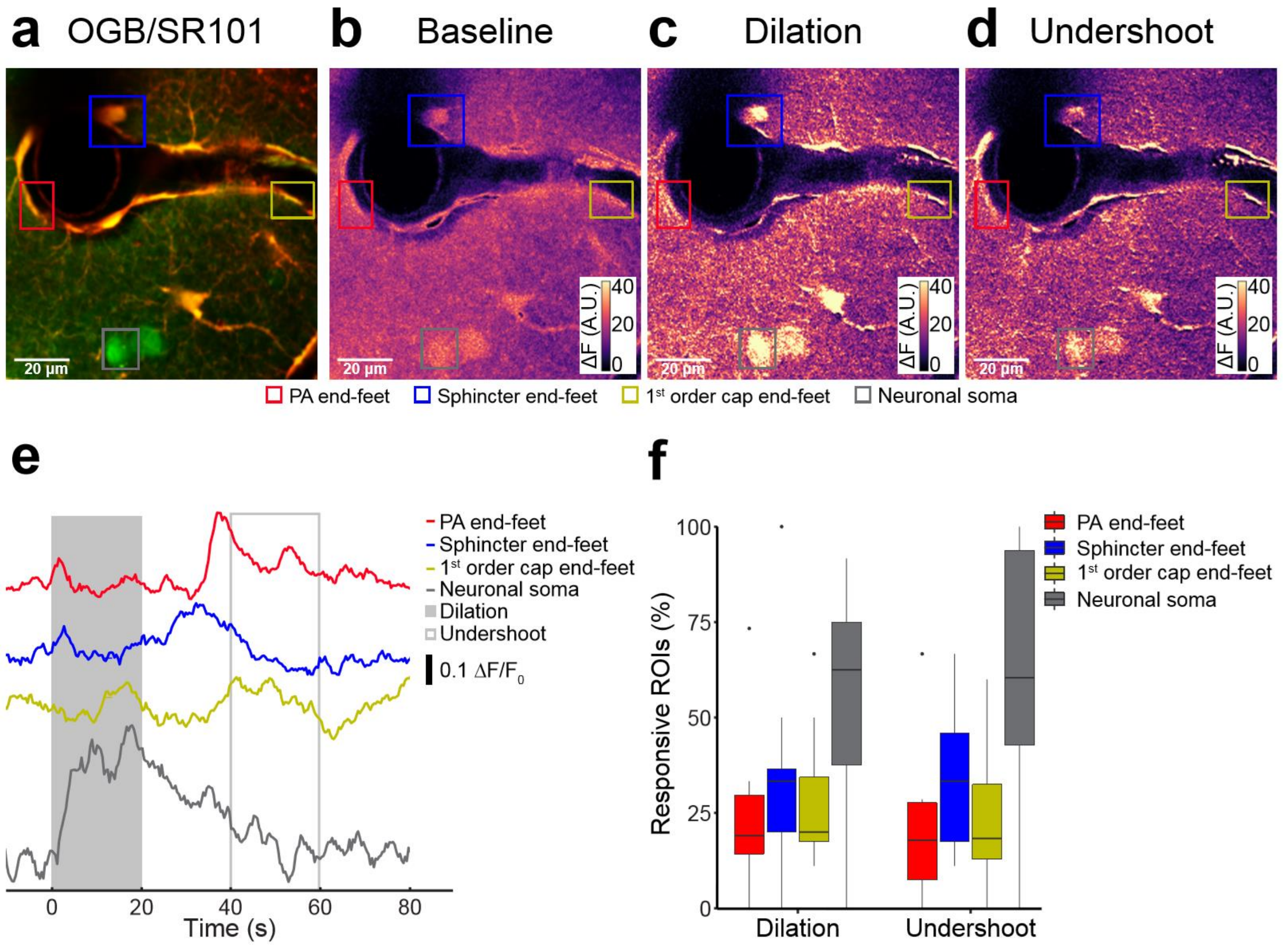


*Extended data Figure 6 | Neuronal somas and astrocytic end-feet increase intracellular $Ca^{2+}$ upon functional stimulation. In vivo whisker pad stimulation experiments (anesthetized C57bl/6j mice) measuring intracellular $Ca^{2+}$ by two-photon microscopy.* ***a****, Representative average projection of a two-photon time-lapse movie. Green indicates Oregon Green 488 BAPTA-1/AM (OGB). Red indicates sulforhodamine 101 (SR101). Colored rectangles indicate the ROIs for intracellular $Ca^{2+}$ measurements for astrocytic end-feet on PA (red), precapillary sphincter (blue), 1st order capillary (yellow), and on neuronal somas (grey).* ***b-d****, OGB is shown in pseudo colors indicating Mean ΔF during baseline (-20-0 s), 2-Hz whisker stimulation (0-20 s) and undershoot (40-60s) phases. ROIs are color coded as in a. Scale bar = 20 µm.* ***e****, Representative smoothed traces of $\Delta F/F_0$ from ROI locations in (****a****) upon stimulation at 2 Hz for 20 s (light grey box) and during the undershoot phase (open light grey box).* ***f****, Boxplot summary of responsive ROIs during dilation and undershoot phase after whisker pad stimulation. We observed increases in intracellular $Ca^{2+}$ in astrocytic end-feet on all vessel segments and in neuronal somas. The fraction of responsive ROIs was similar both during stimulation and in the undershoot phase. We found no difference in the fraction of responsive ROIs between different locations of end-feet on the vascular three. Datasets were analyzed with Paired Wilcoxon signed rank tests to establish difference ($p < 0.05$) between dilation and undershoot phase and between different ROI types.*

#### Extended data methods

*Two-photon imaging of whisker pad stimulation induced evoked $Ca^{2+}$ responses*

Glass micropipettes produced by a pipette puller (P-97, Sutter Instrument) had a resistance of 1.0 ~ 2.0 MΩ. The pipette was loaded with 0.8 mM of the $Ca^{2+}$-sensitive dye Oregon Green 488 BAPTA-1/AM (OGB-AM; Invitrogen) and 0.005 mM of the astrocyte-specific marker sulforhodamine 101 (SR101; Sigma-Aldrich) dissolved in aCSF[2]. The tip of the electrode was inserted into the whisker barrel cortex (layer 2/3). The dye was ejected into the cortex using a pneumatic injector (3–20 PSI, 10–90 s; Pneumatic Pump; World Precision Instruments) guided by the two-photon microscopy. We conducted *in vivo* imaging with two-photon microscopes using a commercial two-photon microscope (FluoView FVMPE-RS, Olympus) equipped with a 25 x1.05 NA-water-immersion objective (Olympus) and a Mai Tai HP Ti:Sapphire laser (Millennia Pro, Spectra Physics). The excitation wavelength was set to 900 nm. The emitted light was filtered to collect red (590-650nm) and green (510-560nm) light from SR101 (astrocytes) and OGB (relative $Ca^{2+}$ changes), respectively. The frame resolution was 0.450 µm/pixel with a 320 x 240 pixels frame and images were taken at a speed of 2.40 frames per second for evoked $Ca^{2+}$ responses.

Evoked $Ca^{2+}$ responses was imaged in one vessel branching from the penetrating arteriole to a first order capillary, including the neck and bulb structure in a single plane.

*Image Analysis of whisker pad stimulation induced evoked $Ca^{2+}$ responses*

In the study of whisker pad stimulation induced evoked $Ca^{2+}$ responses, ROIs were detected using a modification of the pixel-of-interest-based analysis method[3]. ROIs were positioned around astrocytic end-feet and labeled according to their location on PA, sphincter or 1$^{st}$ order capillary. Further ROIs were positioned around neuronal somas and in neuropil. Astrocytic or neuronal structures were recognized based on SR101/OGB staining, cell morphology, and relation to blood vessels[4]. For each frame, we selected pixels showing intensities of 1.5 standard deviations (SD) above the mean intensity of the ROI. The intensities of these pixels were averaged and then normalized to a 15-s baseline period just before stimulation onset, creating a time trace of $\Delta F/F_0$ for every ROI. These time traces were smoothed with a 3-sec moving average to avoid outlier values. A

$Ca^{2+}$ transient was defined as an intensity increase of ≥5% and of ≥2 SD from baseline, having a duration of ≥2.5 s. Recordings were divided into 20-s time bins (dilation and undershoot). Responsivity was defined as the fraction of ROIs with $Ca^{2+}$ transients within time bins per mouse. Recordings lacking evoked $Ca^{2+}$ responses in neuropil were excluded from analysis.

#### Supplementary videos

##### Supplementary video 1 – Thinned skull:

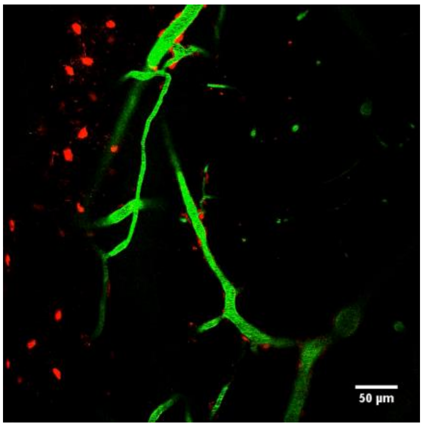


*Supplementary video 1 | Video of a thinned skull in vivo experiment, where the focus shifts in 2µm steps from the skull (notice the NG2-dsRed expression in osteoblasts) through the vasculature in the dura, to the pial arterioles. Closeups of the precapillary sphincters are shown in Extended data Fig. 2.*

##### Supplementary video 2 – Volume rendering of SMA staining:

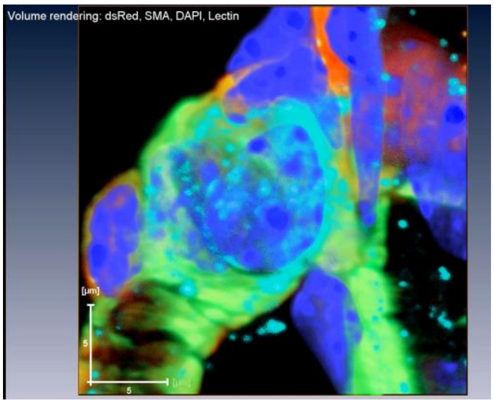


*Supplementary video 2 |Z-stack and volume rendering of a smooth muscle actin antibody and DAPI staining at a precapillary sphincter of an NG2-dsRed mouse brain after i.v. fluorescent-lectin injection.*

#### Supplementary video 3 – 4D whisker pad stimulation:

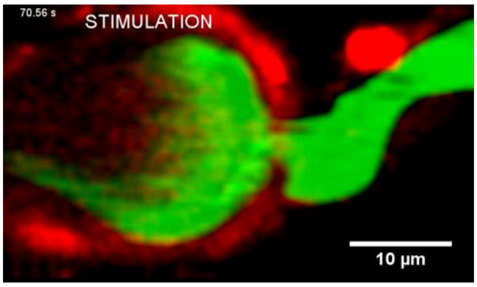

*Supplementary video 3 |Hyperstack projection of a precapillary sphincter dilation after whisker pad stimulation. In the end of the video we show rotation around the x-axis of baseline and stimulation peak.*

#### Supplementary video 4 – RBC stuck at sphincter:

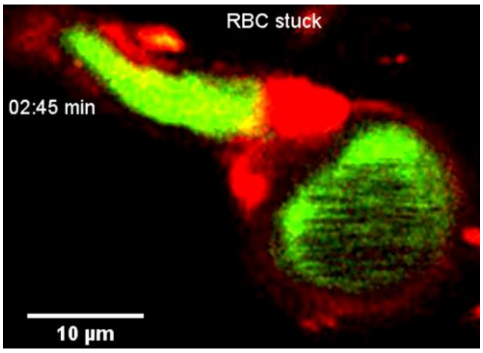

*Supplementary video 4 |Video of RBC's getting stuck temporarily at the precapillary sphincter in an in vivo experiment with a craniotomy (first video) and two pial arteriole precapillary sphincters of a thinned skull experiment (second video).*

#### Supplementary video 5 – RBC deformation

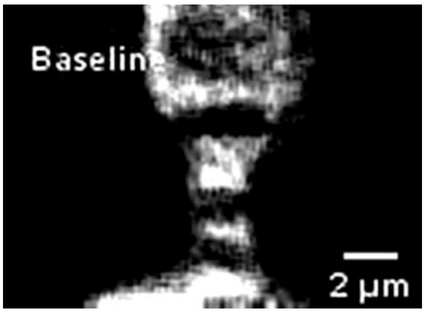

*Supplementary video 5 |Resonance frame-scan video of RBCs passing through a precapillary sphincter and taking up the parachute form. (The RBCs move in the opposite direction of the resonance scan, which will flatten their shapes)*

#### Supplementary video 6 – CSD

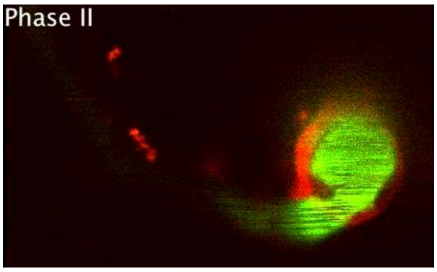

*Supplementary video 6 | Video of the three phases of the CSD in a dsRed mouse with FITC-dextran in the vessel lumen.*

### Supplementary video 7 – Cardiac arrest

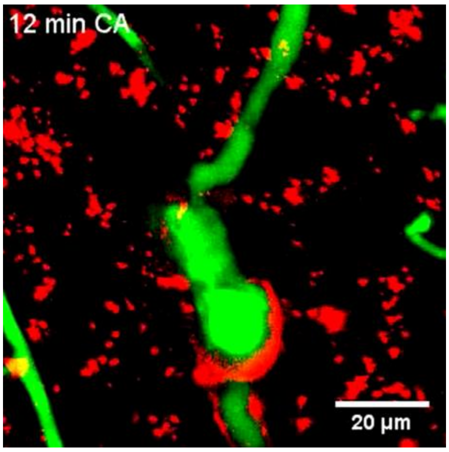

*Supplementary video 7 | Hyperstack recording of two PA branches with precapillary sphincters, in an NG2-dsRed mouse, that collapse in rigor after cardiac arrest (first video). Second video shows an SR101 staining of astrocytes in a FITC loaded WT mouse where the astrocytes swell while the vascular lumen collapses.*

#### Extended data references


1. Silvers, W. K. *The Coat Colors of Mice: A Model for Mammalian Gene Action and Interaction*. (Springer Science & Business Media, 2012).
2. Lind, B. L., Brazhe, A. R., Jessen, S. B., Tan, F. C. C. & Lauritzen, M. J. Rapid stimulus-evoked astrocyte Ca2+ elevations and hemodynamic responses in mouse somatosensory cortex in vivo. *Proc. Natl. Acad. Sci.* **110**, E4678–E4687 (2013).
3. Lind, B. L. *et al.* Fast Ca2+ responses in astrocyte end-feet and neurovascular coupling in mice. *Glia* **66**, 348–358 (2018).
4. Fordsmann, J. C. *et al.* Spontaneous astrocytic Ca2+ activity abounds in electrically suppressed ischemic penumbra of aged mice. *Glia* **67**, 37–52 (2019).
